# Evolutionary dynamics of carbapenem-resistant *Acinetobacter baumannii* circulating in Chilean hospitals

**DOI:** 10.1101/467589

**Authors:** Andrés Opazo-Capurro, Iván San Martín, Mario Quezada-Aguiluz, Felipe Morales, Mariana Domínguez-Yévenes, Celia A. Lima, Fernanda Esposito, Louise Cerdeira, Helia Bello-Toledo, Nilton Lincopan, Gerardo González-Rocha

**Affiliations:** Laboratorio de Investigación en Agentes Antibacterianos (LIAA-UdeC), Departamento de Microbiología, Facultad de Ciencias Biológicas, Universidad de Concepción, Chile; Millennium Nucleus for Collaborative Research on Bacterial Resistance (MICROB-R); Departamento de Farmacia, Facultad de Farmacia, Universidad de Concepción, Chile.; Department of Clinical Analysis, School of Pharmacy, University of São Paulo, São Paulo, Brazil; Department of Microbiology, Institute of Biomedical Sciences, University of São Paulo, São Paulo, Brazil

## Abstract

We analyze the evolutionary dynamics of ninety carbapenem-resistant *Acinetobacter baumannii* (CRAB) isolates collected between 1990 and 2015 in Chile. CRAB were identified at first in an isolate collected in 2005, which harbored the IS*Aba1*-*bla*_OXA-69_ arrangement. Later, OXA-58- and OXA-23-producing *A. baumannii* strains emerged in 2007 and 2009, respectively. This phenomenon was associated with variations in the epidemiology of OXA-type carbapenemases, linked to nosocomial lineages belonging to ST109 (CC1), ST162 (CC79), ST15 (CC15) and ST318 (CC15).

---

Carbapenem-resistant *Acinetobacter baumannii* (CRAB) has been deemed a critical-priority pathogen by the World Health Organization (WHO) (1). It is normally involved in infections acquired in the intensive care units (ICUs), and is commonly resistant to several antibiotics, including carbapenems (2). Accordingly, OXA-type carbapenemases (OTCs) are the main resistance mechanism to carbapenems in *A. baumannii* (3). While OXA-51-like carbapenemases are chromosomally encoded, the remaining OTCs (OXA-23-like, -24-like, -58-like and -143-like) are frequently plasmid encoded (4, 5). OXA-51-like enzymes can mediate resistance to carbapenems if they are overexpressed when the IS*Aba1* element is present upstream of the *bla*_OXA-51-like_ gene (6). CRAB outbreaks are commonly associated to the three predominant clonal complexes (CCs) CC109/1, CC118/2 and CC187/3 (University of Oxford/Institute Pasteur MLST schemes) (7). Although, the clonal complex CC113/CC79 has been predominant in South America; CC104/CC15, CC110/ST25 and CC109/CC1 are also present in this region (8).

The aim of this study was to investigate the evolutionary dynamics of CRAB in Chilean hospitals, where this pathogen has an endemic status.

Ninety non-repetitive *A. baumannii* isolates recovered between 1990 and 2015 were included. They were collected in hospitals from nine different cities throughout Chile, in which the greatest distance between two cities is 2,433 km, representing over 50% of the length of the country.

Antibiotic susceptibility tests were performed to carbapenems, cephalosporins, aminoglycosides, ampicillin/sulbactam, piperacillin/tazobactam, ciprofloxacin, and tetracycline (9). Imipenem (IPM) and meropenem (MEM) MICs were determined following the CLSI guidelines (9). Colistin-resistance was screened using the SuperPolymyxin media (10). Multidrug-resistant (MDR), extensively-drug resistant (XDR) and pandrug-resistant (PDR) phenotypes were defined as previously described (11, 12).

Genetic relatedness was determined by pulsed-field gel electrophoresis (PFGE) as described earlier (13). Groups with at least three genetically related isolates (>87% similarity) were designated as major PFGE clusters (14). Single-locus *bla*_OXA-51-like_ sequence-based typing (SBT) was carried out as described previously (15). Isolates representative of the main PFGE clusters were subjected to whole-genome sequencing (WGS), and sequence types (STs) were determined (Pasteur’s scheme) as published earlier (16).

OTCs genes were screened by multiplex-PCR (17), whereas *bla*_OXA-51-like_ alleles were investigated by PCR and sequencing. IS*Aba1*-*bla*_OXA-51-like_ array was examined by conventional PCR (18). CarbAcinetoNP test was performed on all carbapenem non-susceptible isolates that were negative for *bla*_OXA_ genes (19).

The comprised isolates were grouped into three different periods: P1 (1990-1999, *n*= 27), P2 (2000-2009, *n*= 30), and P3 (2010-2015, *n*= 33). Consequently, carbapenem resistance was confirmed in 56 (62%) isolates, being identified for the first time in 2005 in a strain (A329, P2) carrying the IS*Aba1*-*bla*_OXA-69_ array (Figure 1). XDR, MDR or PDR profiles were displayed by 51 (57%), 28 (31%) and 3 (3%) isolates, respectively. Furthermore, 65 (72%) isolates were non-susceptible to amikacin, whereas 64 (71%) were non-susceptible to gentamicin. Additionally, 32 (36%) isolates exhibited resistance to ampicillin-sulbactam, and 4 (3.6%) were colistin-resistant.

**Figure 1.**
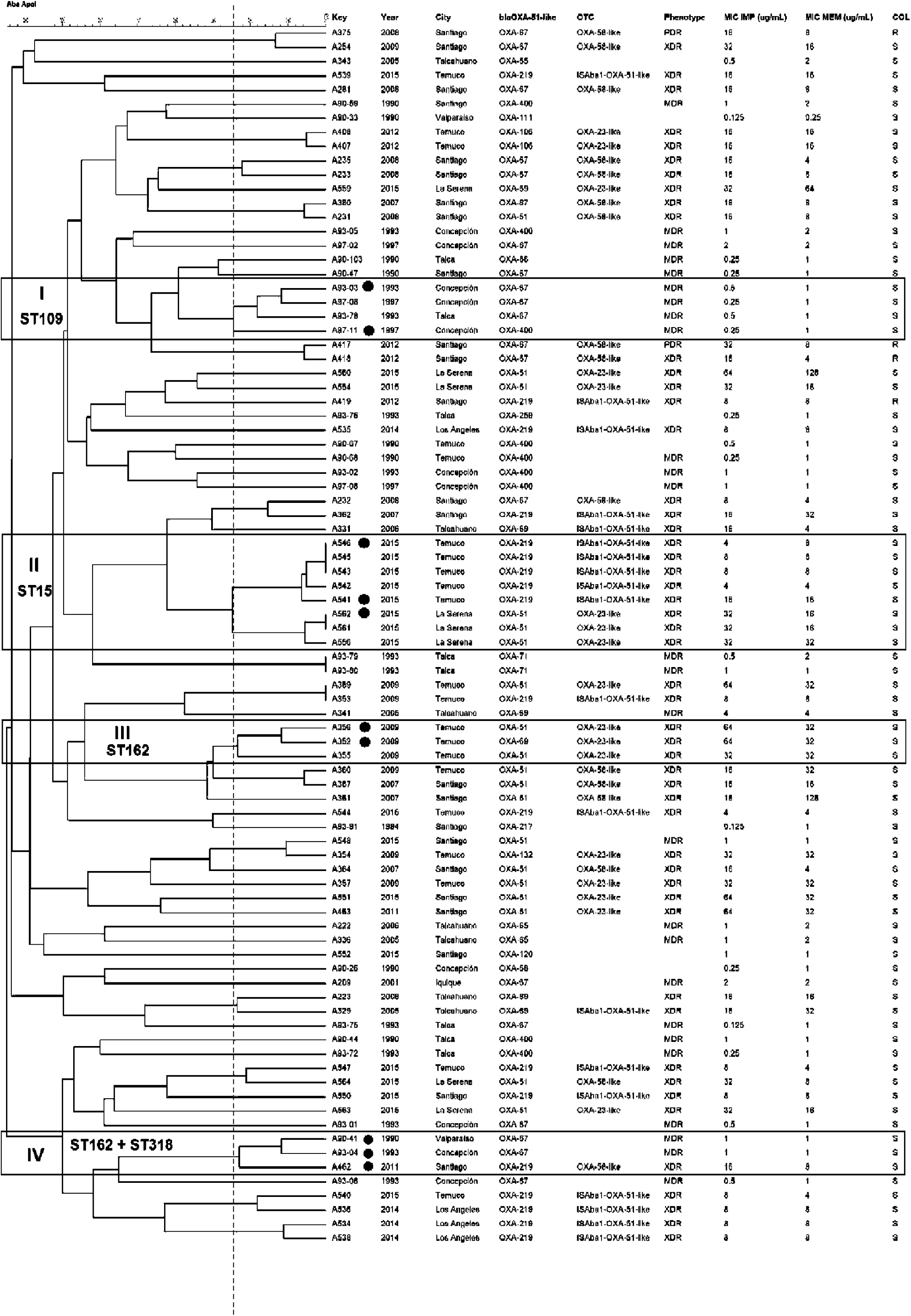
Dendrogram generated after restriction with ApaI enzyme for 86/90 typed *A. baumannii* isolates. The black dotted line represents 87% similarity. I to IV denote the major PFGE groups characterized according to the criteria described in the manuscript. MDR: multidrug-resistant; XDR: extensively-drug resistant; PDR: pandrug-resistant; OTC: OXA-type carbapenemase; ST: Sequence type; COL: colistin. ● Isolates typified by MLST.

Further, *bla*_OXA-58_ (30%) and *bla*_OXA-23_ (30%) genes were more prevalent and were associated with highest carbapenems MICs (Figure 1). The IS*Aba1*-*bla*_OXA-219_ array was observed in 14 of 56 (25%) CRAB isolates. In this regard, OXA-58-producing isolates seems to have emerged in 2007, whereas IS*Aba1*-OXA-219 and OXA-23 producers arose in 2009, being disseminated among different hospitals.

As expected, no OTC producers were identified in P1. Otherwise, eleven OXA-58-, seven OXA-23-, and four IS*Aba1*-*bla*_OXA-51_-like-positive CRAB isolates were detected in P2 (Figure 1). In P3, a change in the molecular epidemiology of circulating OTCs was observed, where OXA-23 producers (*n*= 11) were predominant, followed by OXA-51-like (associated with ISAba1, *n*= 15)- and OXA-58 (*n*= 4)-positive isolates (Figure 1). Interestingly, the CRAB isolate A223 was negative for both CarbAcinetoNP and OTCs PCR. Thus carbapenem-resistance could be mediated by a different mechanism (2).

Four major clusters (I – IV) were identified by PFGE (Figure 1). Cluster I included four carbapenem-susceptible isolates from P1, while cluster II comprised CRABs from 2015 that harbored the OXA-23-like (*n*=3) and IS*Aba1*-*bla*_OXA-219_ array (*n*=5), which were collected from two hospitals separated by >1000 km (Figure 1). Cluster III contained three CRABs carrying *bla*_OXA-23_-Hke genes and the OXA-51-like variants OXA-51 and OXA-69. Finally, cluster IV included three isolates from three different cities, comprising a single OXA-58-like-producing CRAB (Figure 1). Four isolates were non-typeable.

Fifteen *bla*_OXA-51_-like variants were identified from SBT, where most prevalent alleles were OXA-51 (*n*= 21), OXA-67 (*n*= 20) and OXA-219 (*n*= 18) (Figure 1). They are not associated to the three predominant international clones (ICs). Furthermore, isolates from PFGE cluster I corresponded to ST109, whereas those from clusters II and III belonged to ST15 and ST162, respectively (Figure 1). In cluster IV, two isolates from P1 belonged to ST109, whereas a single isolate (A462) from P3 corresponded to ST318, which is part of the CC15.

In Chile, CRAB has been responsible for about 26% of ventilator-associated pneumonia (VAP) in hospitalized adults (20), whereas carbapenem-resistance rates are above 66% (21). Our results reveal the evolutionary dynamics of CRAB in the country, focusing on the major carbapenem resistance genes and lineages circulating in hospital settings in a period of 25 years.

Worryingly, XDR isolates were predominant in our collection, including resistance to aminoglycosides and ampicillin/sulbactam, in concordance with previous reports in the country (22). Although the rate of colistin resistance was 3.6%, this percentage is higher than the previously published in 2012 (1.4%) (22), representing an alarming increase to be considered CRAB has been increasing lately worldwide, and our results reveal that initially in Chile it was related to the IS*Aba1*-*bla*_OXA-69_ array identified in 2005, where ISs play an essential role in the regulation of this resistance (23).

Concerning to acquired OTCs, OXA-58-like-producing isolates seem to have emerged in 2007, whereas OXA-23-like producers arose later (3, 24). Significantly, after 2010 a new change in the molecular epidemiology of circulating OTCs was observed, where OXA-23 producers have been predominant and widely disseminated along the country. Additionally, we detected the replacement of certain carbapenem-susceptible clones present in P1, by carbapenem-resistant linages that began to emerge in the late 2000s. SBT revealed that the CRAB isolates were not related to the major ICs (I-III). The main OXA-51-like variants present were OXA-219, OXA-67 and OXA-51. Of these, OXA-51 has been associated with the CC15 (15), previously detected in Europe, Pakistan and South America, which is considered as a high-risk clone (25). In South America, this CC is categorized as epidemic in Brazil (26), which suggests the dissemination of resistant clones through the region. Otherwise, OXA-67 and OXA-219 are related to less prevalent ICs (15). Interestingly, OXA-219 was originally identified in 2012 from a single isolate from Chile, being related to the worldwide (WW) clone 4 (27), associated to the IS*Aba1*-*bla*_OXA-219_ array. These results suggest the presence of an endemic lineage (WW4, OXA-219) coexisting with a regional lineage (ST15) in Chile (8, 28), which has been described in Brazil (29) and Ecuador (28).

Other identified lineages included ST109 (CC1), ST162 (CC79), and ST318 (CC15). ST109 has been originally identified in Sweden (30), whereas ST162 and ST318 have been described in Brazil (29, 31). These findings reaffirm that the major lineages present in the region are different to those globally spread (8). However, ICII and III have been lately identified in Peru (32), which might have an important impact on the local epidemiology.

In conclusion, our study provides data about evolutionary dynamics of CRAB circulating in Chilean hospitals, which were linked to particular lineages as well as to the emergence of specific OTCs, whereas colistin resistance deserves an urgent attention to strengthen surveillance.

## Acknowledgments

The authors want to thank to the National Fund for Scientific and Technological Development (FONDECYT) for supporting this study. The authors also want to thank the microbiologists of the hospitals and medical centers who kindly provided the isolates including in this project. N.L. is a research grant fellow of Conselho Nacional de Desenvolvimento Cientifico (CNPq 312249/2017-9). This work was partially presented at the American Society for Microbiology (ASM) Microbe 2016 meeting, Boston, USA.

## Conflict of Interest Statement

None to declare.

## Funding Source

This work was supported through funds granted by the National Fund for Scientific and Technological Development (FONDECYT) of Chile (project N°3150286), the Universidad de Concepción (project ENLACE-VRID N°216.036.044-1.0), Fundação de Amparo à Pesquisa do Estado de São Paulo, Brazil (FAPESP 2016/08593-9) and Conselho Nacional de Desenvolvimento Científico, Brazil (CNPq 462042/2014-6).

